# *easybio*: an R Package for Single-Cell Annotation with CellMarker2.0

**DOI:** 10.1101/2024.09.14.609619

**Authors:** Wei Cui

## Abstract

Single-cell RNA sequencing (scRNA-seq) allows researchers to study biological activities at the cellular level, enabling the discovery of new cell types and the analysis of intercellular interactions. However, annotating cell types in scRNA-seq data is a crucial and time-consuming process, with its quality significantly influencing downstream analyses. Accurate identification of potential cell types provides valuable insights for discovering new cell populations or identifying novel markers for known cells, which may be utilized in future research. While various methods exist for single-cell annotation, one of the most common approaches is to use known cell markers. The CellMarker2.0 database, a human-curated repository of cell markers extracted from published articles, is widely used for this purpose. However, it currently offers only a web-based tool for usage, which can be inconvenient when integrating with workflows like Seurat. To address this limitation, we introduce *easybio*, an R package designed to streamline single-cell annotation using the CellMarker2.0 database in conjunction with Seurat. *easybio* provides a suite of functions for querying the CellMarker2.0 database locally, offering insights into potential cell types for each cluster. In addition to single-cell annotation, the package also supports various bioinformatics workflows, including RNA-seq analysis, making it a versatile tool for transcriptomic research.

## 1. Introduction

Accurately identifying the specific cell types in single-cell data is critical for many downstream analyses. Numerous methods have been developed for single-cell annotation.

Recent research has evaluated the annotation accuracy of several methods, including GPT-4, SingleR, and CellMarker2.0 [1]. The SingleR method [2], a supervised approach, relies on reference datasets for accuracy and can be time-consuming. scType [3] integrates databases like PanglaoDB [4] and the original CellMarker database [5] to annotate single cells using both positive and negative markers. However, CellMarker has been updated to version 2.0 [6], which incorporates new marker resources and is manually curated for human and mouse cell types.

While direct use of CellMarker2.0 may not always achieve the highest annotation accuracy, it offers broad applicability across diverse datasets and provides more interpretable results. However, its functionality has so far been limited to an online user interface, with no software implementation available [1].

To address this, we introduce the R package *easybio* [7], designed to provide convenient access to the CellMarker2.0 database for marker querying and streamlined single-cell annotation.

## 2. R package *easybio*

### 2.1. Query Markers in the CellMarker2.0 Database

One of the key features of the CellMarker2.0 database [6] is its ability to search for markers based on the top differentially expressed genes in each cluster, helping to determine the potential cell type for each cluster. The package also includes functions that allow users to retrieve markers along with information about their corresponding tissues, as reported in published studies. The package also supports retrieving markers directly for specific cell types.

**Figure 1:**
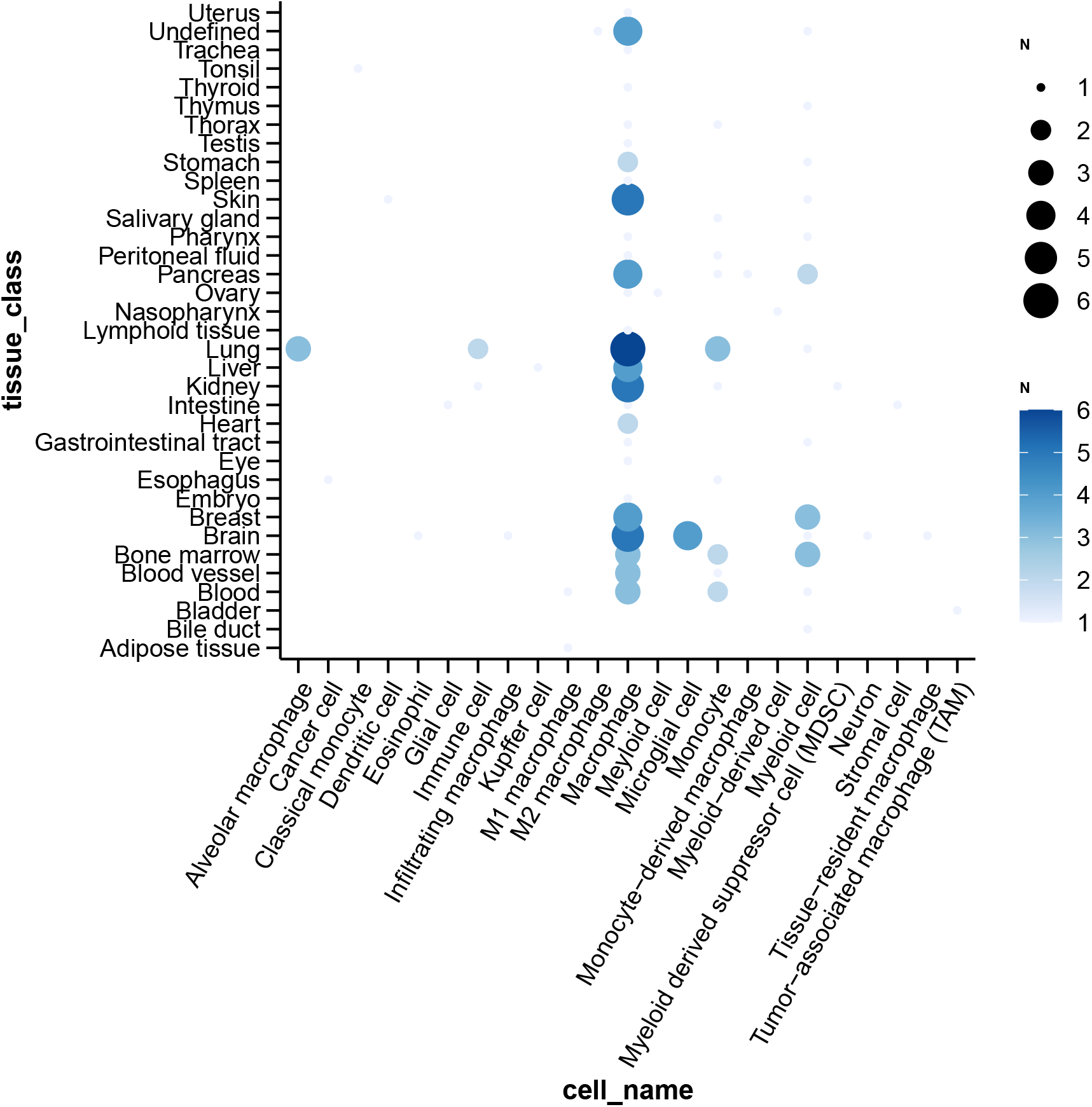
Using the R package *easybio*, a query for CD68 in CellMarker2.0 illustrates the distribution of this marker across different tissues and cell types

### 2.2. Annotate Single-Cell Clusters with CellMarker2.0

Annotating cell clusters is a crucial step in single-cell RNA-seq analysis, helping to assign biological identities to each group of cells. Typically, this process involves comparing the differentially expressed genes (DEGs) between clusters and identifying the most enriched genes within a cluster. These enriched genes serve as markers that can be used to determine the potential cell types within each cluster.

The CellMarker2.0 database is a valuable resource for this purpose, as it provides a curated collection of cell-type markers derived from published studies. While the web-based tool allows researchers to manually search for markers by pasting gene lists, this approach can be time-consuming and may require matching one cluster at a time. For large datasets, this manual process can introduce inefficiencies and limit the speed of analysis.

To overcome these limitations, our package automates the process by directly matching the top genes from each cluster to potential cell types using the CellMarker2.0 database. This not only speeds up the annotation process but also reduces the likelihood of manual errors. The package also provides flexibility by allowing users to specify the number of top-ranked genes (n) used for matching, enabling finer control over the annotation process. This feature is particularly useful for optimizing the balance between marker specificity and sensitivity.

While it may be tempting to adopt the top-matched cell type as the definitive annotation for each cluster, we encourage users to explore additional matched cell types. In cases where multiple, distinct cell types are matched to the same cluster, it is important to investigate the biological context and the broader experimental conditions. This exploratory approach can help uncover novel or rare cell populations and ensure that the annotation is both accurate and comprehensive. By leveraging the full potential of CellMarker2.0, users can enhance the resolution of their single-cell analysis and gain deeper insights into cellular heterogeneity.

## 3. Example Workflow

In this example workflow, we use the PBMC3K dataset and R package Seurat [8] to illustrate the process.

### 3.1. Execute the Seurat PBMC3K Guided Tutorial

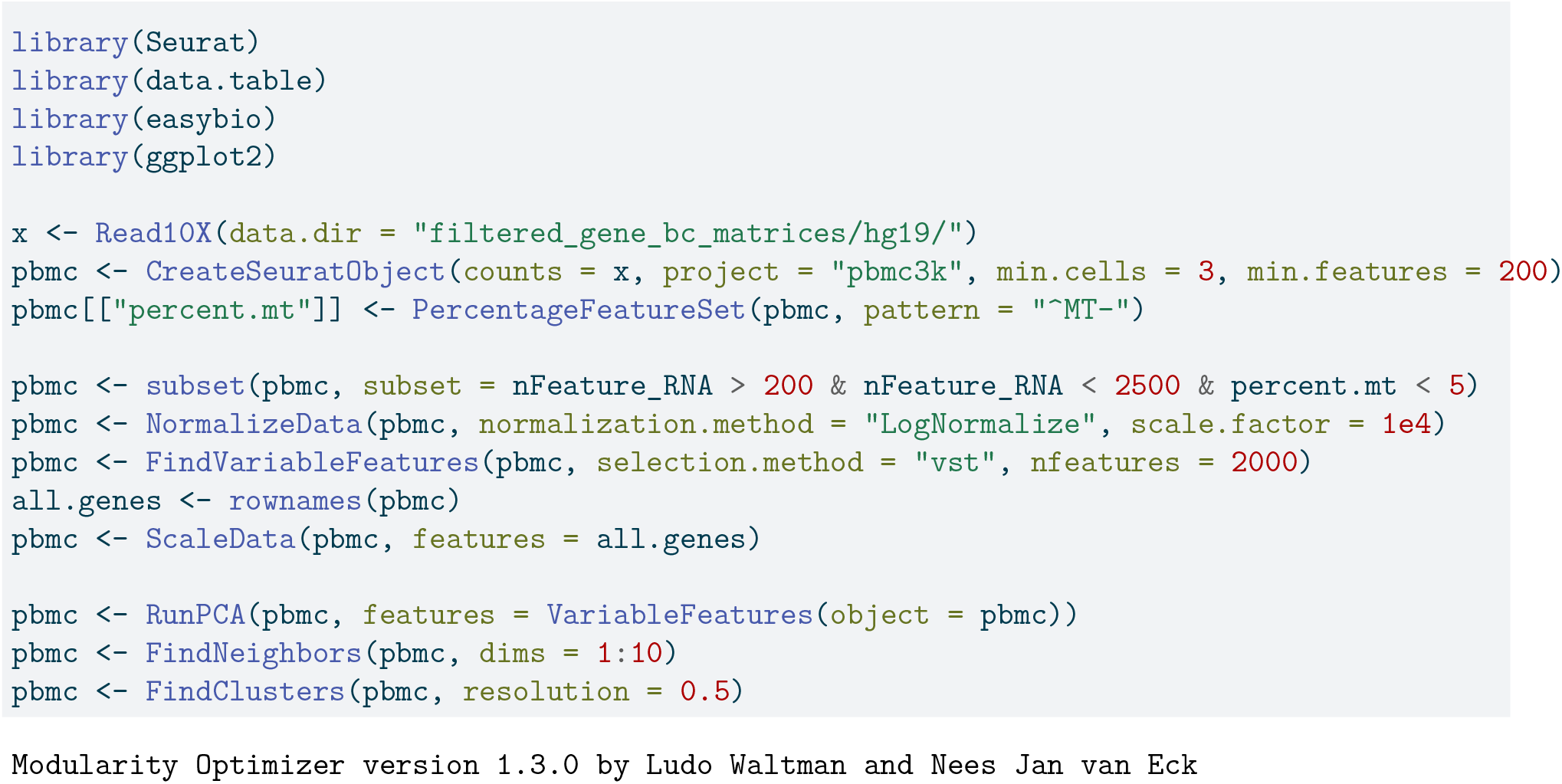
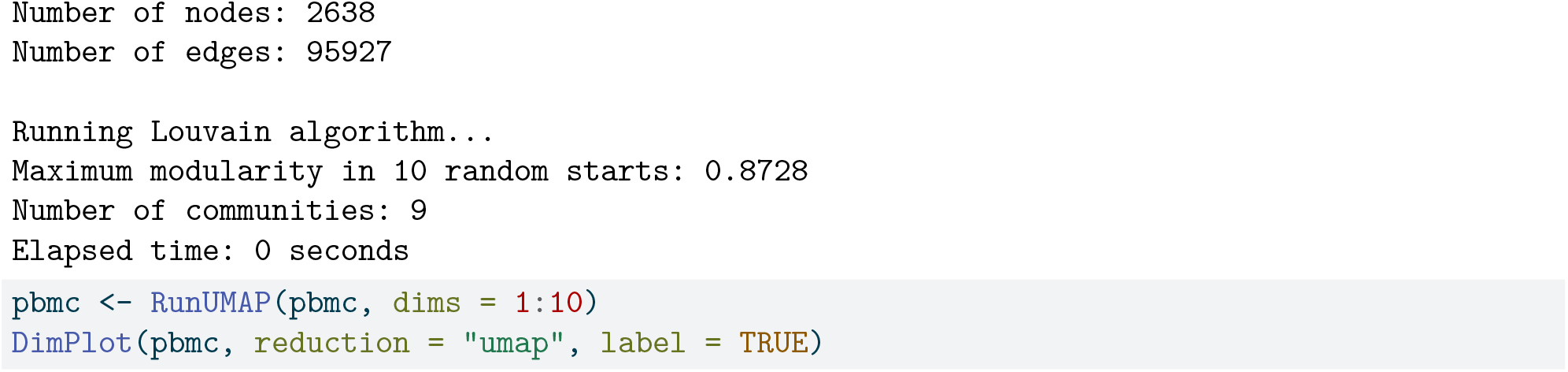

**Figure 2:**
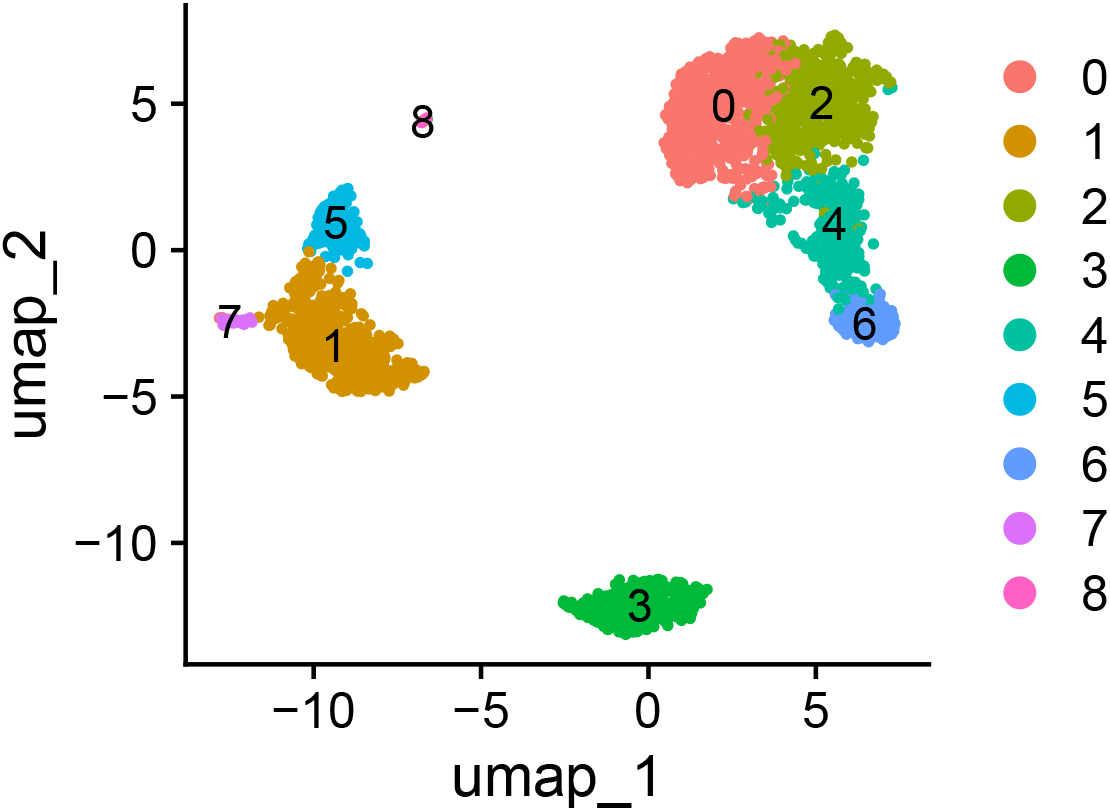
UMAP of raw, unannotated Clusters

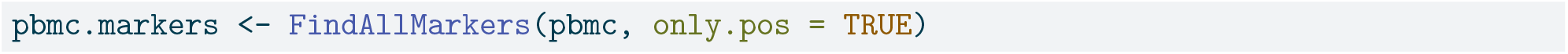

### 3.2. Match with CellMarker2.0

For each cell cluster, we identify the top 50 most highly expressed genes (selecting only those with an adjusted p-value < 0.05 and ordering them by decreasing log2 fold change) and use these to query the CellMarker2.0 database for matching cell type markers. This approach allows us to align the gene expression profiles with known markers, facilitating cell type annotation.

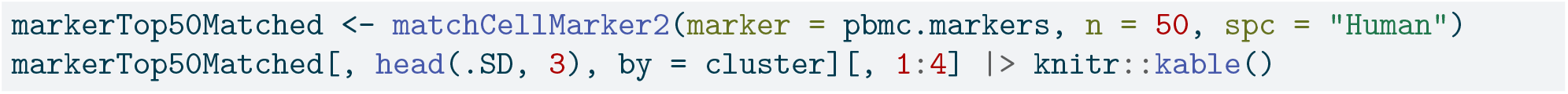

**Table 1:**
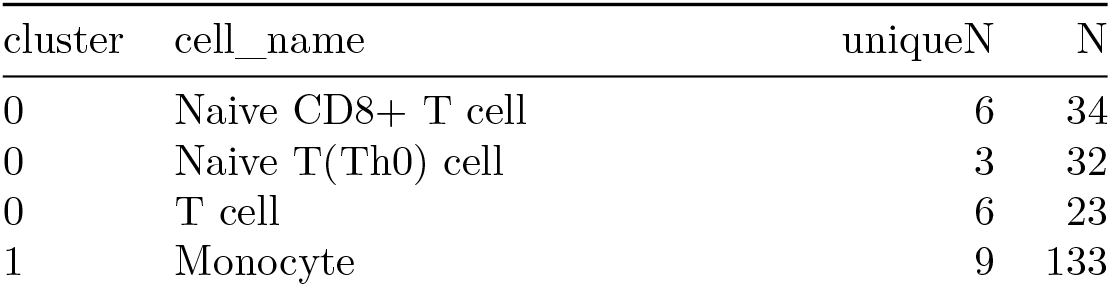
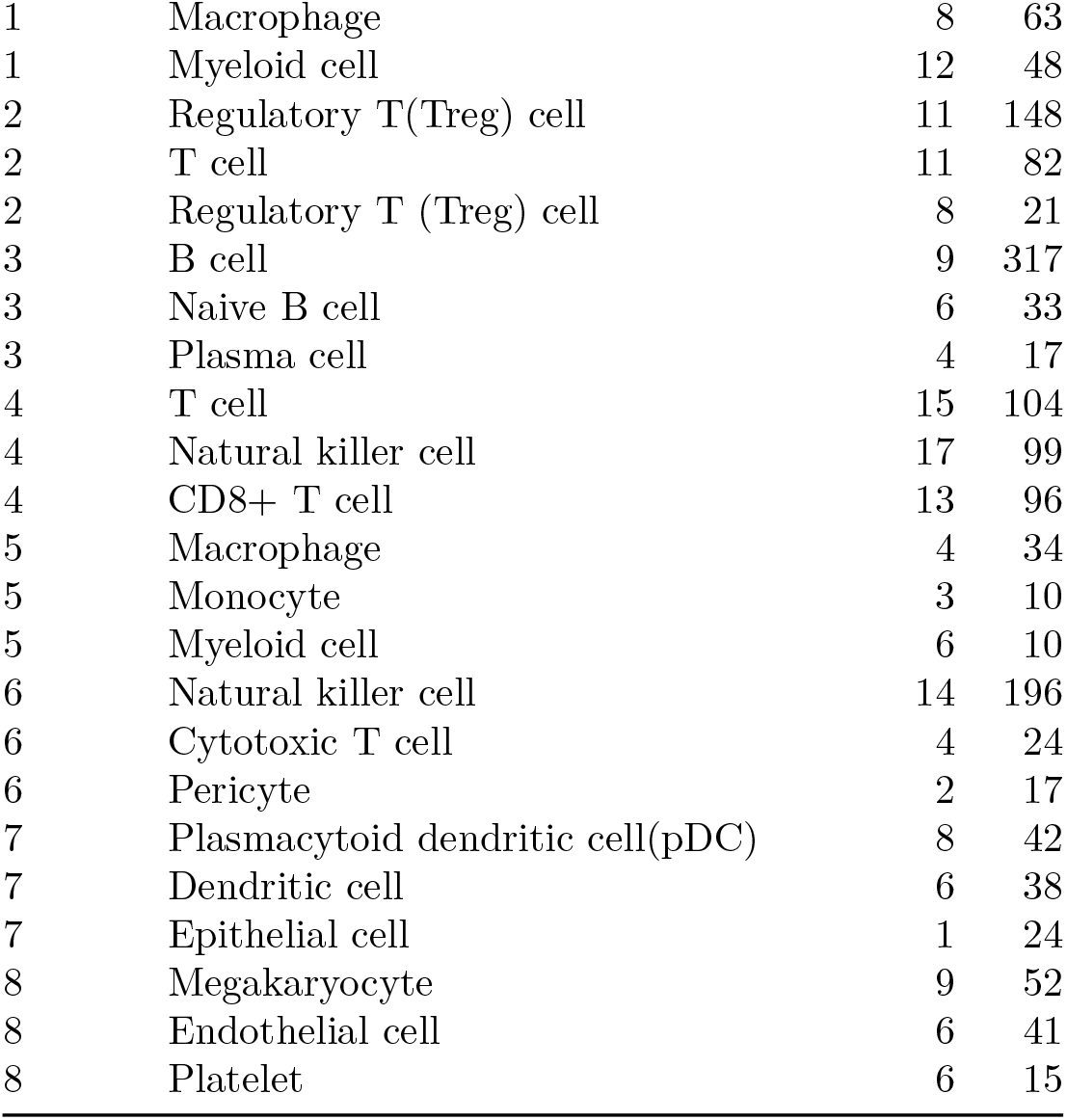
The matched marker numbers for each cluster in the CellMarker2.0 database are shown. The column labeled “N” represents the total number of matched markers, while “uniqueN” indicates the number of unique markers. Additionally, the count of each marker is also recorded.

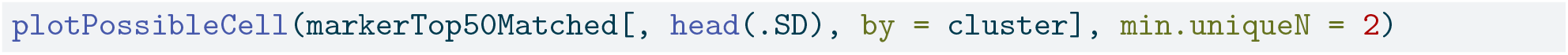

**Figure 3:**
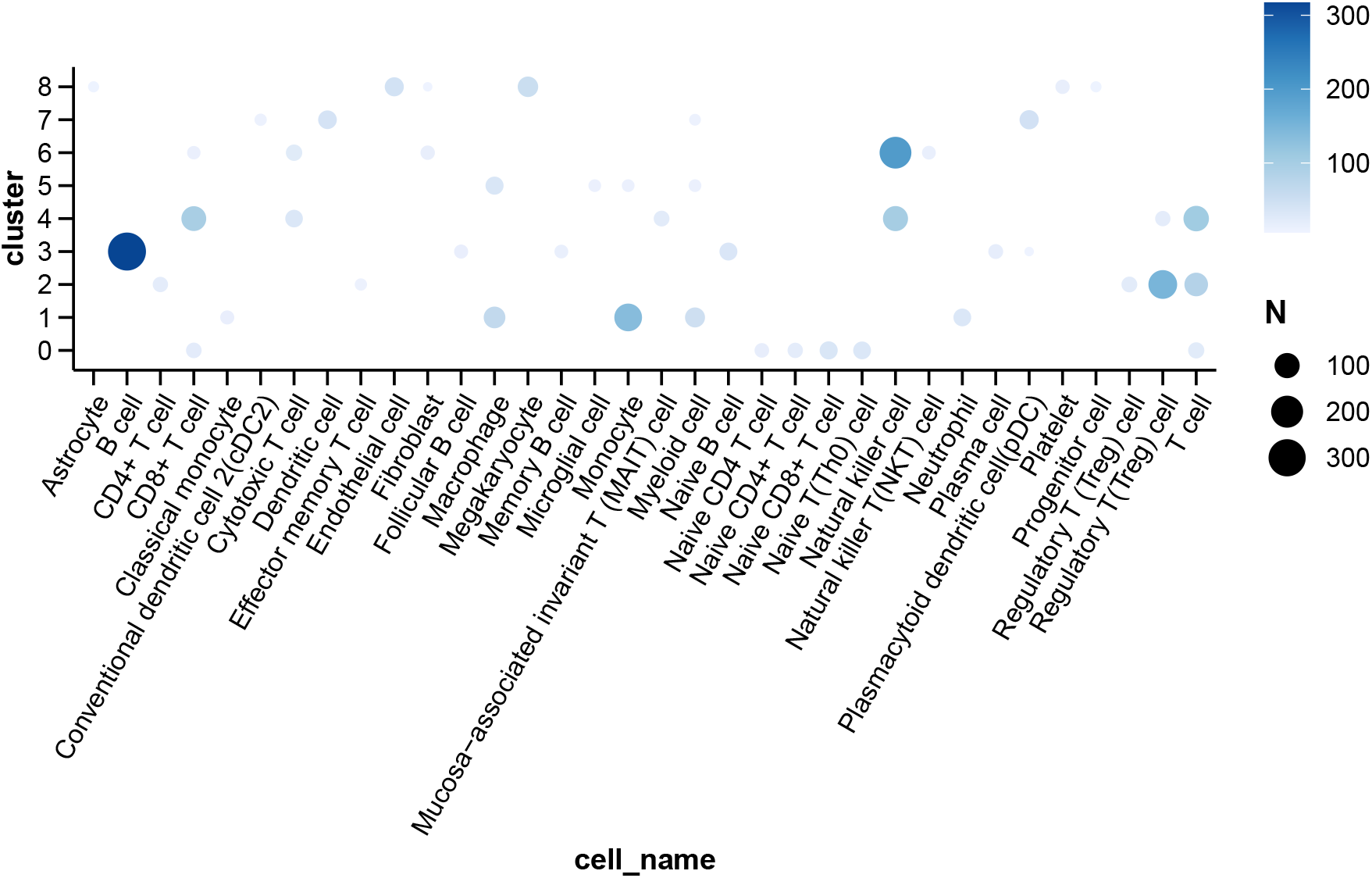
Visualization of cell clusters and their corresponding cell types

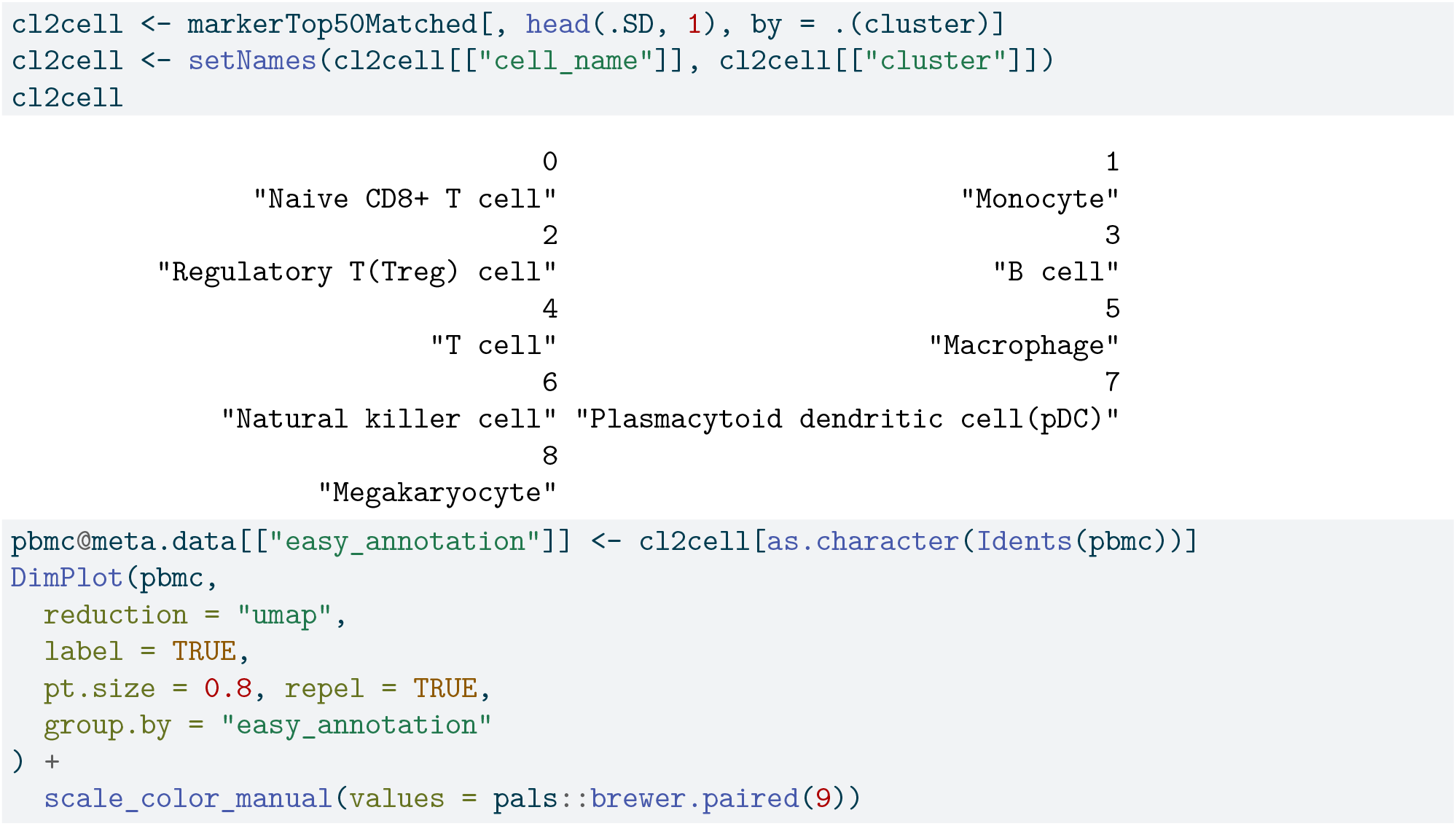

**Figure 4:**
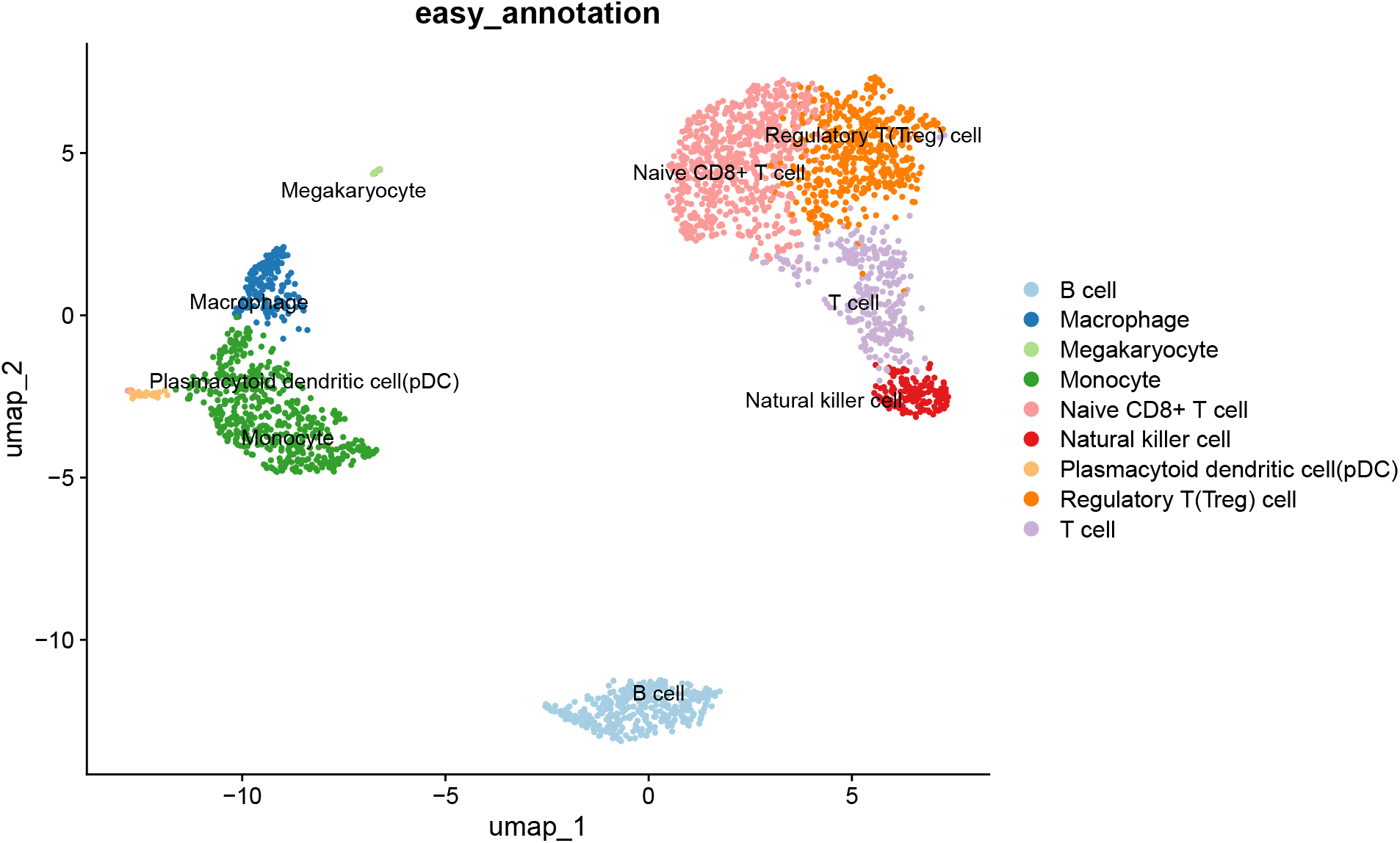
UMAP of easy annotation using the top matched cell.

### 3.3. Evaluate Additional Potential Cell Types

Although the top-matched cell type is commonly used for annotation, it is recommended to also review the expression of markers for other potential cell types. This is particularly important when a cluster is matched to several distinct cell types. Such evaluation helps to ensure more accurate and reliable annotations. To streamline the process, clusters that are close together in the UMAP plot, such as clusters 1, 5, and 7, can be examined simultaneously.

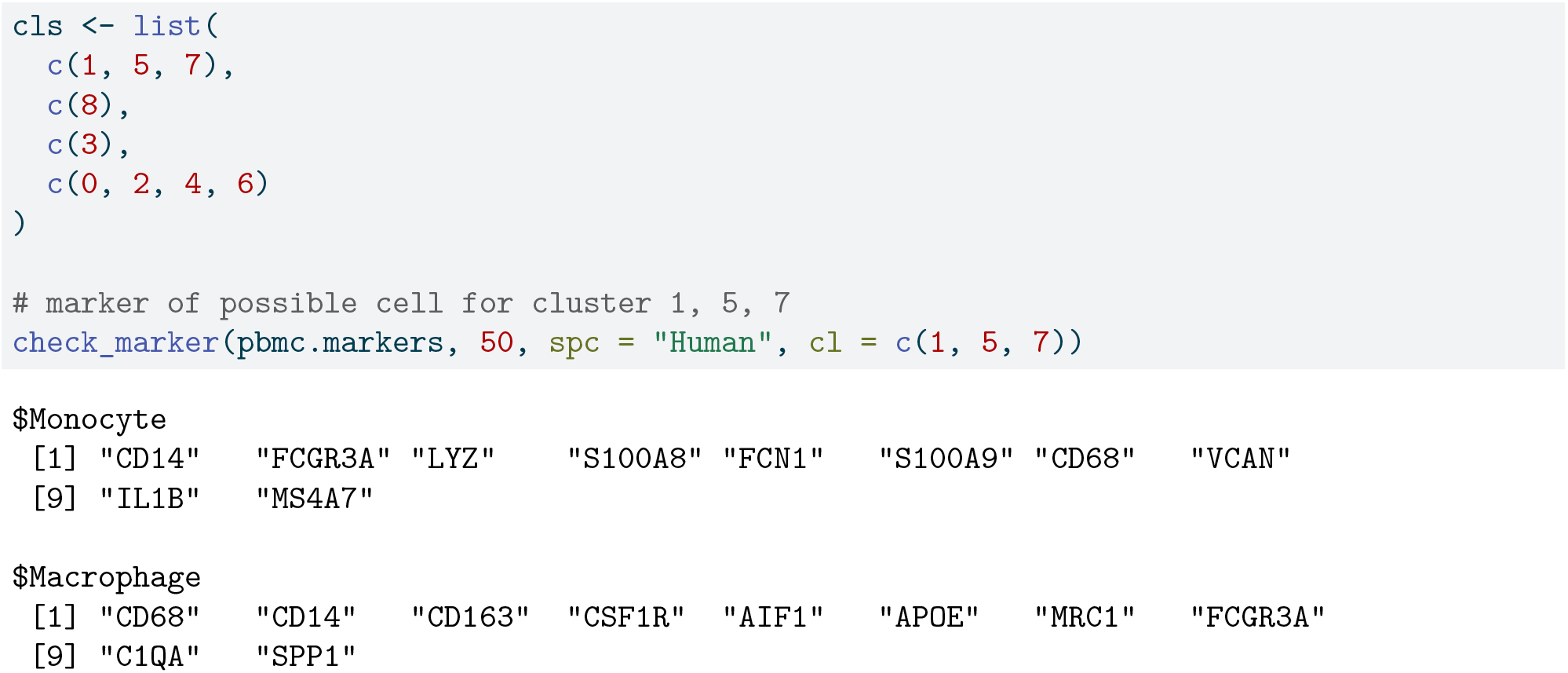
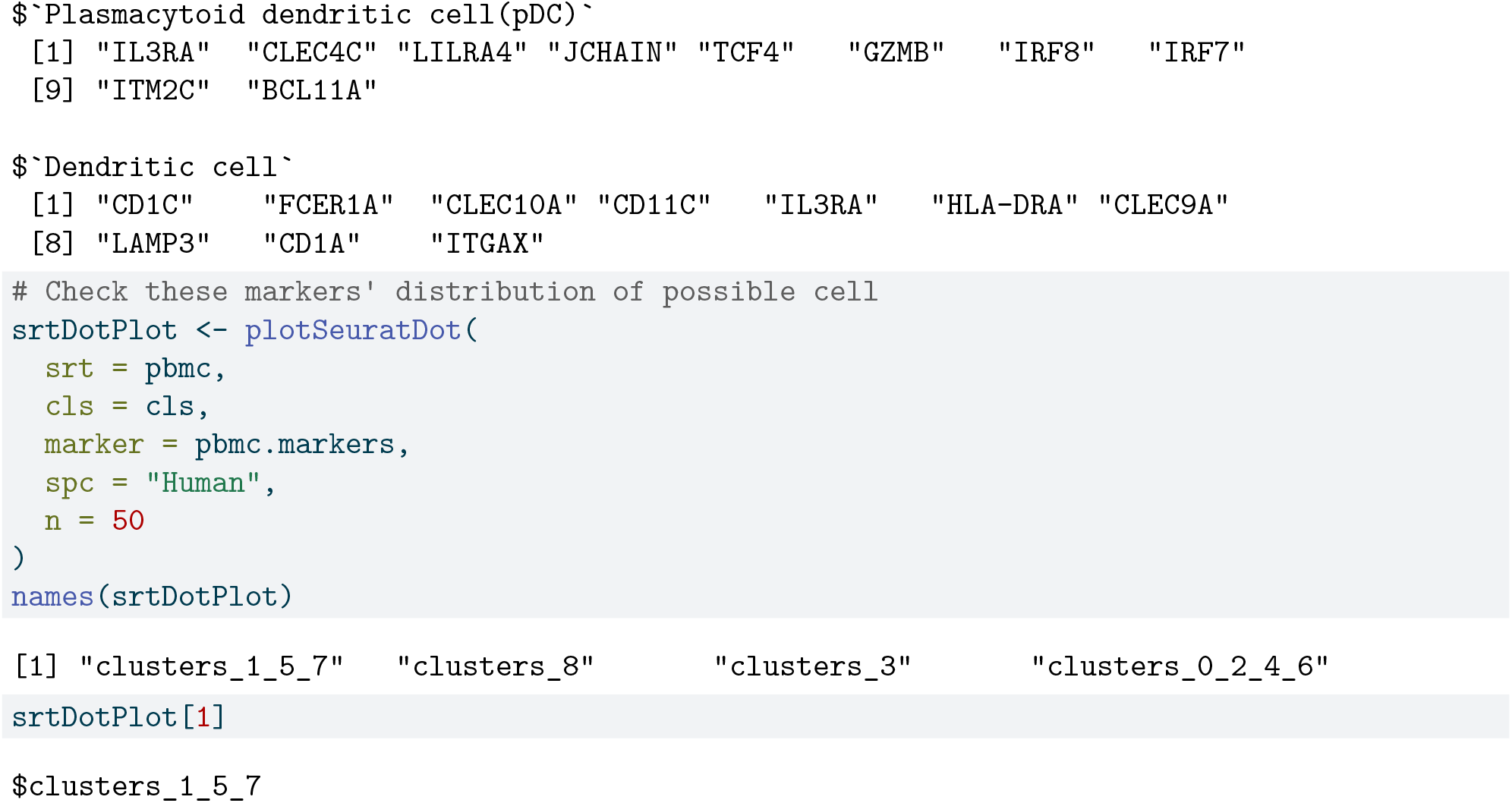

**Figure 5:**
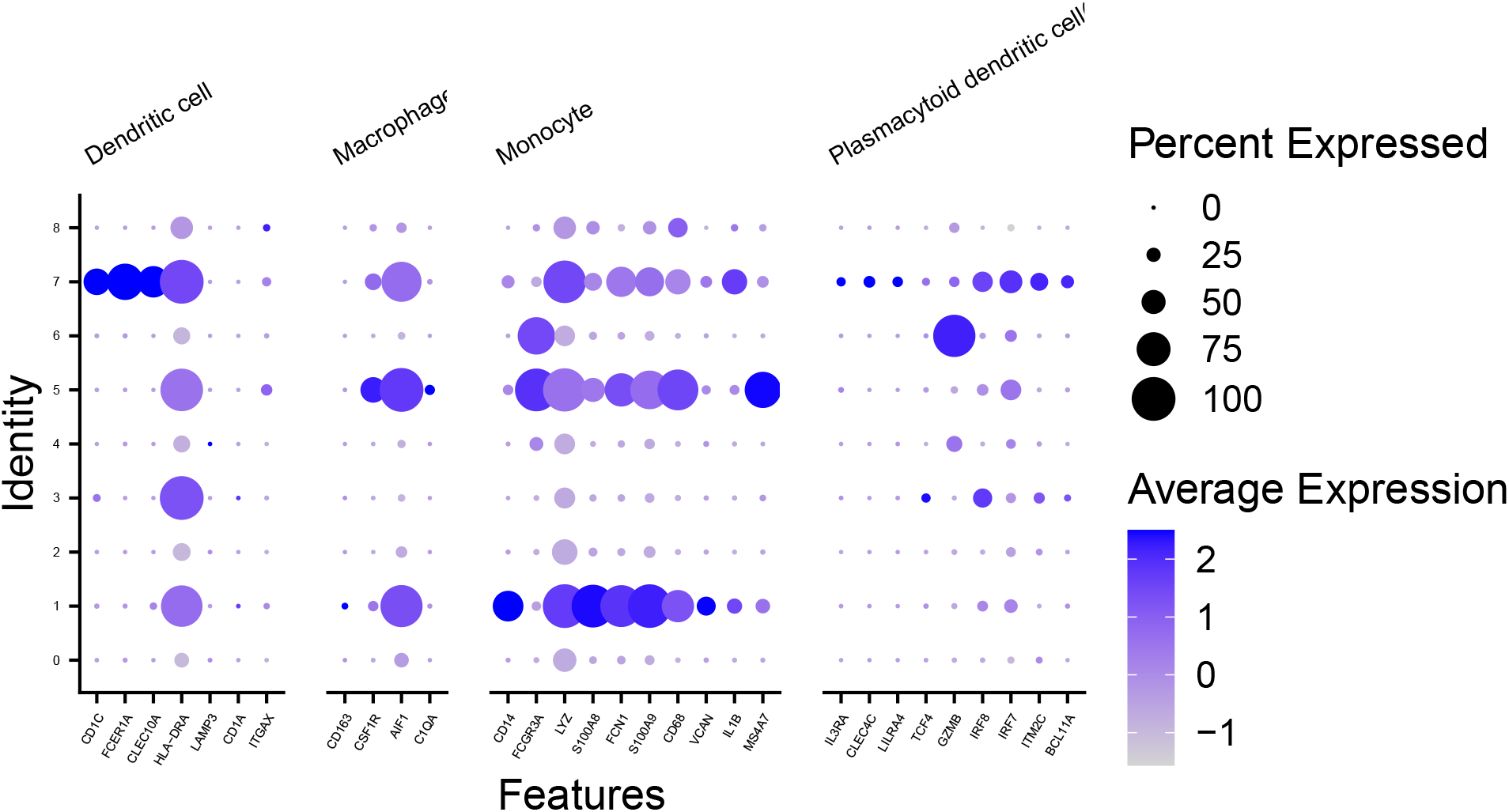
Marker expressions for potential cell types are shown for clusters 1, 5, and 7. Expressions for other clusters are omitted for brevity.

Based on the dot plot, create a named vector to assist with cell annotation.

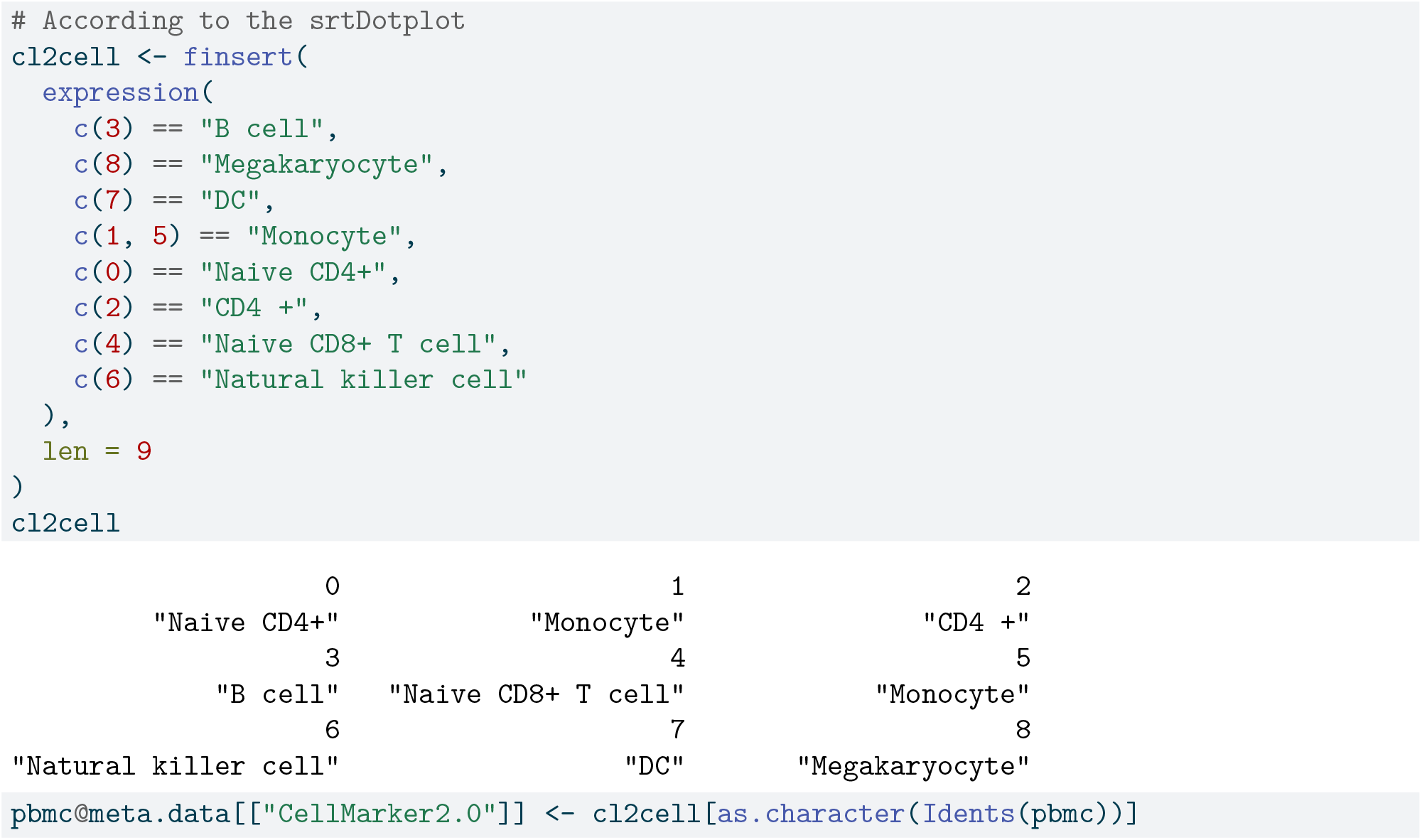

### 3.4. Compare with SingleR

In this analysis, we use the widely-adopted R package SingleR [2] to annotate the data. This allows us to compare the annotation results obtained from CellMarker2.0 with those generated by SingleR, providing a basis for evaluating the accuracy and consistency of our annotations.

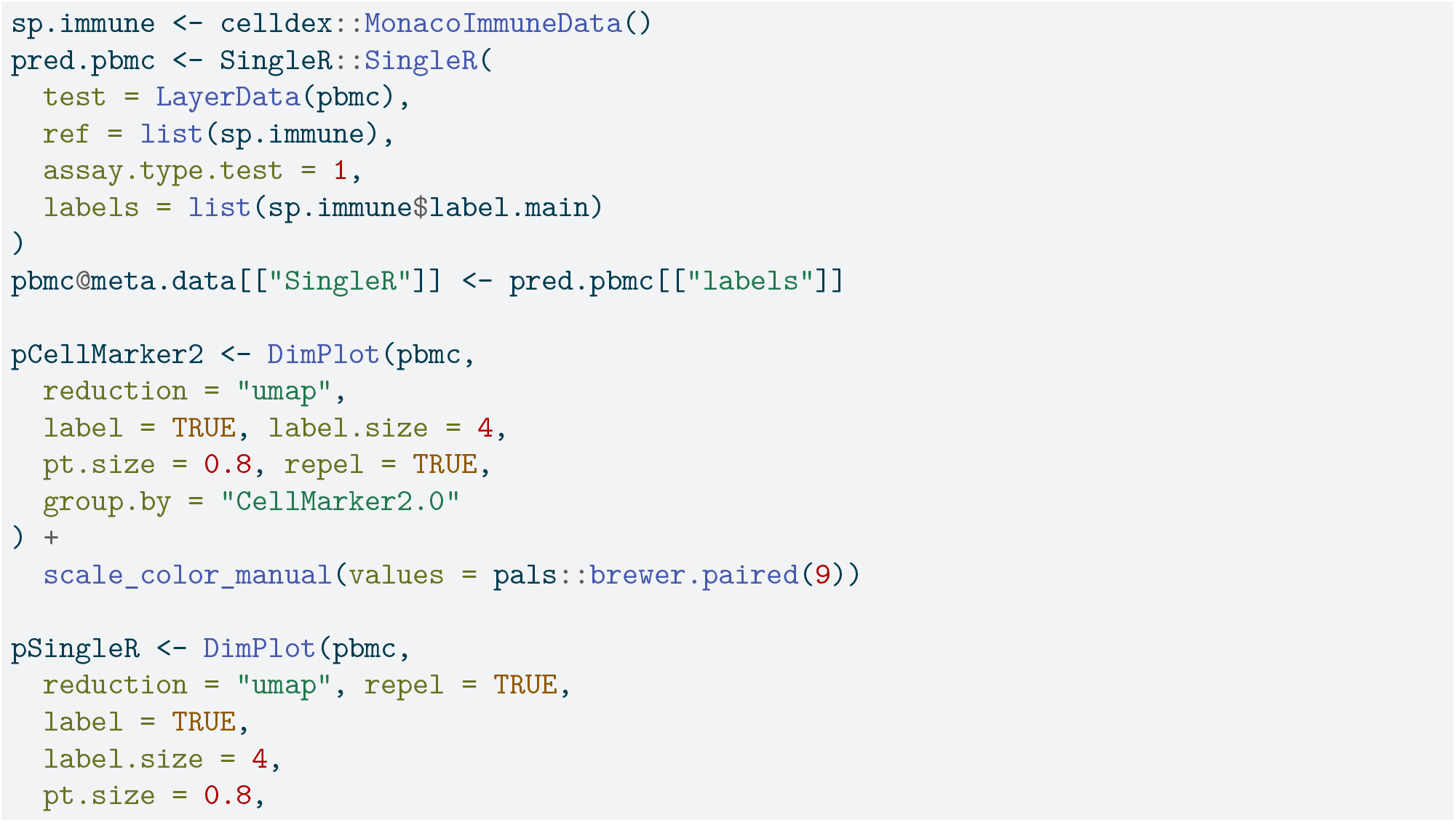
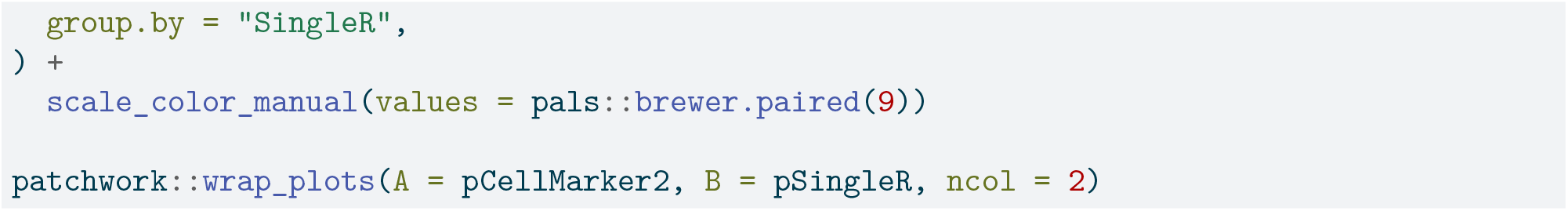

**Figure 6:**
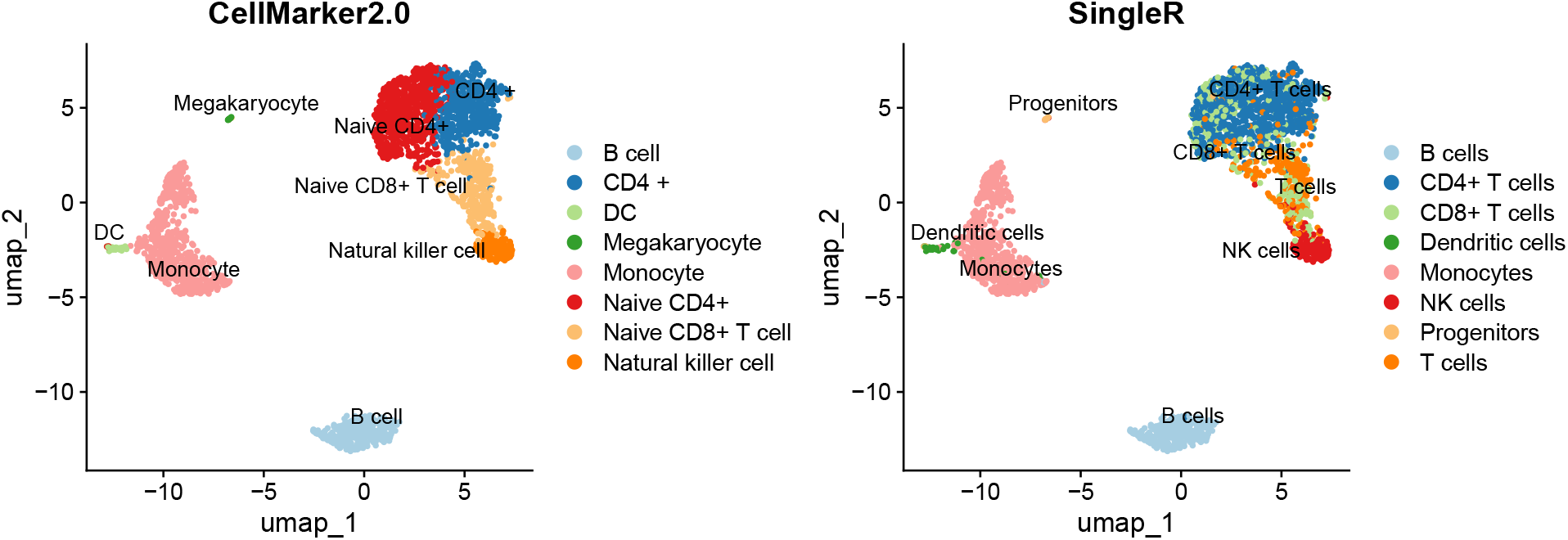
Comparation of annotation from CellMarker2.0 and SingleR

## 4. Conclusion and Discussion

In this study, we introduced the R package *easybio*, designed to facilitate single-cell annotation by leveraging the CellMarker2.0 database. To our knowledge, *easybio* is the first R package to incorporate CellMarker2.0 for this purpose.

We assessed the package’s performance by applying it to the Seurat pbm3k tutorial dataset and comparing the resulting annotations with those obtained using the SingleR package and manual annotations through Seurat. The comparison demonstrated that annotations derived from CellMarker2.0 were consistent with those from SingleR and Seurat’s manual methods. Importantly, *easybio* operates independently of external reference datasets, thereby reducing the time and expertise required compared to manual annotation processes.

*easybio* is not limited to single-cell annotation with CellMarker2.0; it also supports a range of analyses including bulk RNA-seq and data exploration, and it facilitates integration with other databases.

Nonetheless, several limitations should be acknowledged. The effectiveness of single-cell annotation with CellMarker2.0 depends on the quality of cell clustering, which is influenced by preprocessing steps such as data quality assessment, principal component analysis (PCA), and the selection of resolution parameters. Variations in these parameters can lead to different clustering outcomes and, consequently, affect annotation results. It is recommended to experiment with different parameter settings to assess their impact. Additionally, we only evaluated performance using the pbmc3k dataset. A broader range of datasets should be tested to provide a more comprehensive assessment, and we did not employ standardized methods to rigorously evaluate accuracy.

In summary, *easybio* simplifies the single-cell annotation workflow by incorporating the CellMarker2.0 database, offering a more efficient and reproducible tool for researchers.

## 5. Data availability

The R package is now available on The Comprehensive R Archive Network (CRAN). The single-cell marker files can be downloaded directly from CellMarker2.0. The PBMC3K data can be accessed here.

## 6. Acknowledgements

Special thanks to the amazing R package ggplot2 [9] for providing an easily extensible grammar of graphics, and to data.table [10] for its concise syntax and ultra-fast data processing capabilities. We also appreciate the journal template and ChatGPT for assisting with language refinement.

## 7. Author contributions statement

WC conceived and designed the study, conducted the experiments, and analysed the results.

## References

[1] W. Hou, Z. Ji, Assessing GPT-4 for cell type annotation in single-cell RNA-seq analysis, Nature Methods 21 (8) (2024) 1462–1465, publisher: Nature Publishing Group. doi:10.1038/s41592-024-02235-4. URL https://www.nature.com/articles/s41592-024-02235-4

[2] D. Aran, SingleR: Reference-Based Single-Cell RNA-Seq Annotation, r package version 2.6.0 (2024). doi:10.18129/B9.bioc.SingleR. URL https://bioconductor.org/packages/SingleR

[3] Fully-automated and ultra-fast cell-type identification using specific marker combinations from single-cell transcriptomic data | Nature Communications. URL https://www.nature.com/articles/s41467-022-28803-w

[4] O. Franzén, L.-M. Gan, J. L. M. Björkegren, PanglaoDB: a web server for exploration of mouse and human single-cell RNA sequencing data, Database: The Journal of Biological Databases and Curation 2019 (2019) baz046. doi: 10.1093/database/baz046. URL https://www.ncbi.nlm.nih.gov/pmc/articles/PMC6450036/

[5] X. Zhang, Y. Lan, J. Xu, F. Quan, E. Zhao, C. Deng, T. Luo, L. Xu, G. Liao, M. Yan, Y. Ping, F. Li, A. Shi, J. Bai, T. Zhao, X. Li, Y. Xiao, CellMarker: a manually curated resource of cell markers in human and mouse, Nucleic Acids Research 47 (Database issue) (2019) D721–D728. doi:10.1093/nar/gky900. URL https://www.ncbi.nlm.nih.gov/pmc/articles/PMC6323899/

[6] C. Hu, T. Li, Y. Xu, X. Zhang, F. Li, J. Bai, J. Chen, W. Jiang, K. Yang, Q. Ou, X. Li, P. Wang, Y. Zhang, CellMarker 2.0: an updated database of manually curated cell markers in human/mouse and web tools based on scRNA-seq data, Nucleic Acids Research 51 (D1) (2022) D870–D876. doi:10.1093/nar/gkac947. URL https://www.ncbi.nlm.nih.gov/pmc/articles/PMC9825416/

[7] W. Cui, easybio: Comprehensive Single-Cell Annotation and Transcriptomic Analysis Toolkit, r package version 1.0.1 (2024). URL https://github.com/person-c/easybio

[8] R. Satija, Seurat: Tools for Single Cell Genomics, r package version 5.1.0 (2024). URL https://satijalab.org/seurat

[9] H. Wickham, ggplot2: Elegant Graphics for Data Analysis, Springer-Verlag New York, 2016. URL https://ggplot2.tidyverse.org

[10] T. Barrett, M. Dowle, A. Srinivasan, J. Gorecki, M. Chirico, T. Hocking, B. Schwendinger, data.table: Extension of ‘data.frame’, r package version 1.16.99 (2024). URL https://r-datatable.com

